# The first glimpse of *Homo sapiens* hereditary fusion genes

**DOI:** 10.1101/2022.05.06.490969

**Authors:** Degen Zhuo

## Abstract

Family-inherited fusion genes have been known to be associated with human disease for decades. However, only a small number of them have been discovered so far. In this report, monozygotic (MZ) twins are used as a genetic model to investigate hereditary fusion genes (HFG). We have analyzed RNA-Seq from 37 MZ twins and discovered 1,180 HFGs, the maximum of which is 608 per haploid genome. Eight HFGs associated with MZ twin inheritance range from 52.7% to 67.6%, some of which are previously-studied cancer fusion genes and indicate hereditary cancer genes. These data suggest that HFGs are major genetic factors for human diseases and complex traits. This study gives us the first glimpse of human HFGs and lays theoretical and technological foundations for future genetic and medical studies.

---

A gene was thought to be a unit of inheritance that ferried a characteristic from parent to child^1^. Fusion genes such as *BCR-ABL1* had been traditionally thought to be somatic and cancerous^2,3^ and, hence, not hereditable^4^. Human family-inherited fusion genes generated by genomic alterations were responsible for significant inherited pathology of humans (Homo sapiens)^5–7^. Human germline genomic structural variants (SV) were the genetic foundation of hereditable fusion genes^8^. Limitations available to genome technologies historically hindered accurate SV identification^9,10^. As genome technologies progressed from array-comparative genomic hybridization to long-read sequencing and other emerging technologies, the prevalence of human genomic SVs had dramatically increased from about 300 to 34,234 SVs per human haploid genome^10^. However, these complex SVs were often mapped to multiple locations in a genome, which made it impossible to obtain reproducible data for genetic studies^7^. During the last several decades, traditional molecular cloning has been extensively used for investigating SV-associated human diseases, such as human color blindness, which was discovered in 1998^11^; inherited peripheral neuropathies^12^; and other diseases^12–16^.

Recent advances in RNA-Seq technologies have identified large numbers of fusion transcripts, most of which were thought to be somatic and cancerous^17^. On the other hand, many studies have shown that many fusion transcripts exist in high frequencies in non-cancer tissues^18–20^. In addition to read-through fusion transcripts^20^, fusion transcripts resulting from genomic alterations were first discovered in cancer but later in healthy samples. *TPM4-KLF2^21^*, *PIM3-SCO2^22,23^, NCO2-UBC^24^*, and *OAZ1-KLF2^21^* were first reported in cancer samples, the first three of which were later observed in normal controls^24^. These apparent contradictions suggest that fusion genes require further exploitation. Recently, RNA-Seq has been used to identify fusion transcripts associated with rare inherited diseases^25^. Previously, we developed SCIF (splicing codes identify fusion gene transcripts) to more accurately and efficiently discover fusion transcripts from RNA-Seq datasets and identified enormous numbers of fusion transcripts^26^. However, when we systematically validated cancer-specific fusion genes such as *KANSARL* (*KANSL1-ARL17A*), they were detected in healthy samples and individuals at very high frequencies^26^. Eventually, *the KANSARL* gene was validated as the predisposition (hereditary) fusion genes^26^. To study family-inherited fusion genes more precisely, we defined the hereditary fusion gene (HFG) as the fusion gene that offspring inherited from parents and excluded read-through fusion transcripts generated via transcriptional termination failure. Since environmental and physiological factors regulated read-through^20,27,28^, we defined the epigenetic fusion gene (EFG) as the fusion genes generated via *cis*-splicing of read-through pre-mRNAs of two same-strand neighbor genes of the human reference genome. The main differences between HFGs and EFGs were that HFGs existed in ≤50% of human populations while the genes to generate EFGs were present in ≥99.99% of the human population. This report used monozygotic (MZ) twins, who share identical genetic materials^29^, as a genetic model to study human HFGs systematically.

Since MZ twins shared identical genetic materials and even identical epigenetics^29^, Fig.1a showed that an identical HFG (indicated by ‘H’) was carried by a fertilized egg and inherited by two identical embryos. If a random SV mutation (indicated by ‘S’ in Fig.1a) generated a somatic fusion gene, it would be detectable only in one of the MZ twin siblings^30^. If a random SV mutation to generate a fusion gene per individual had a rate of 3.6×10^-2 31,32^, the probability that two identical MZ twins had a random somatic fusion gene was 1.3×10 The probability that both individuals of an MZ twin had an identical HFG was 1/37 or 2.7% and 20-fold higher than that of the random somatic mutations. We used SCIF to analyze blood RNA-Seq data from 37 pairs of MZ twins (dbGap accession: phs000886) and identified 97,770 fusion transcripts. From these fusion transcripts, we identified a total of 1180 HFGs, shown in Supplementary Table 1, in both siblings of ≥1 pair of MZ twins (bHFG), whose frequencies range from 1 to 23 pairs of MZ twins (Fig.1b). These MZ 1180 bHFGs counted only 1.2% of the total fusion transcripts and 15.2% of 7,750 fusion transcripts detected in ≥2 individuals (Supplementary Table 1), suggesting that the MZ bHFGs were not due to random chances. Two hundred seventy-one (23%) of 1180 HFGs had been observed in ≥2 pairs of MZ twins (Fig.1b), the average of which was 3.97 pairs of MZ twins. In addition, Fig.1c showed that 946 (80.2%) of bHFGs had been found to have 1–18 HFGs present in one of two MZ twin siblings (iHFG), the average of which was 4.88 iHFGs. If a bHFG existed in ≥1 pair of MZ twins, the chance of its iHFG generated via a random ‘S’ mutation was ≤3X10^-4^, which was at least 46 fold less than the observed iHFG frequencies, which ranged from 1.4% to 25%. Therefore, iHFG was equal to its counterpart of the bHFG and would be treated as the HFG unless specified. To get the total numbers of HFGs each individual had, we added bHFGs and iHFGs together (Supplementary Table 2). On average, each of 1180 HFGs was detected in 7.3 persons or 9.86% of the 74 MZ twin individuals, which was statically significantly higher than the ‘S’ random mutation^31,32^. Therefore, random somatic ‘S’ rearrangements were mathematically impossible to generate HFGs among healthy populations under no selective environments.

**Figure 1.**
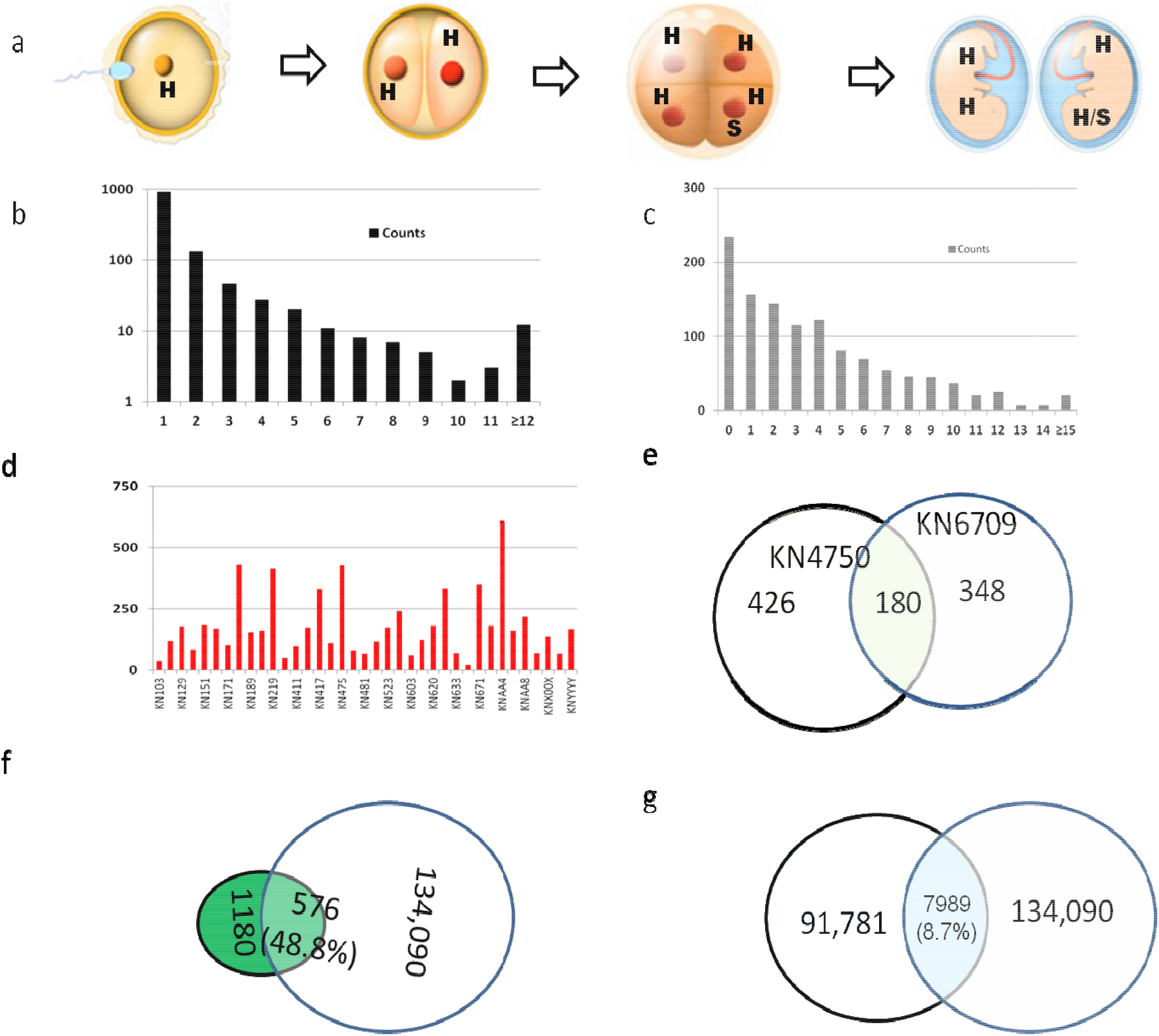
Brief review and characterization of identification of hereditary fusion genes (HFG). a). schematic diagrams to show the formation of two monozygotic (MZ) twin siblings from a monozygote. ‘H’ and ‘S’ represented the hereditary fusion gene and a random somatic genomic alteration. b). recurrent gene frequencies of bHFGs among 37 pairs of MZ twins. bHFGs were fusion genes detected in both siblings of an MZ twin. c). recurrent gene frequencies of iHFGs among 1180 HFGs. iHFGs were fusion genes detected in one individual of an MZ twin among 1180 HFGs. e) Venn diagram showing overlapped HFGs between KN6790 and KN4750 MZ twins. Light blue and white circles represented KN4750 and KN6790 MZ twins, respectively. f) Venn diagram displayed overlapped MZ HFGs and the fusion transcripts discovered in 427 GTEx blood samples. Dark green and white circles showed MZ HFG and the fusion transcripts discovered in 427 GTEx blood samples. g). Venn diagram showed the overlapped numbers of fusion transcripts between 37 pairs of MZ twins and 427 GTEx blood samples. Light blue and white circles showed MZ HFG and the fusion transcripts discovered in 427 GTEx blood samples.

Next, we analyzed the HFG distribution among the MZ twins’ individuals. Fig.1d showed that MZ twins ranged from 21 HFGs in KN650 MZ twins to 608 HFGs in KNAA4 MZ twins. The average MZ twin individual encoded 178.8 HFGs (Supplementary Table 2). Since we showed previously that numbers of fusion transcripts correlated with high-quality RNA-Seq data and RNA-Seq data sizes^26^, the enormous differences among different MZ twin individuals may be mainly due to RNA-Seq qualities. Supplementary Table 3 showed that KN650 and KNAA4 MZ twins had similar RNA-Seq data sizes. KNAA4 MZ twins, SRR2105729 and SRR2105730, had 6212 and 7153 fusion transcripts from which 498 and 510 HFGs, respectively, were identified, while KN650 MZ twins, SRR2105720 and SRR2105721, had 82 and 163 fusion transcripts from which three and 18 HFGs, respectively, were found (Supplementary Table 4). Therefore, the actual number of HFGs would be significantly higher, suggesting that HFGs were widespread and highly diverse among different individuals.

To demonstrate HFG complexities, we selected and compared KN4750 and KN6709 MZ twins. Supplementary Table 2 showed that KN4750 and KN6709 had 426 HFGs and 348 HFGs. A comparison of KN4750 and KN6709 showed that the 180 HFGs overlapped and accounted for 51.7% of KN4750’s 348 HFGs and 42.3% of KN6790’s 426 HFGs (Fig.1e). To generate the overlapping 180 HFGs, KN4750 and KN6709 were expected to have 673 and 1007 HFGs, respectively. KN6709 who had potential 1007 HFGs confirmed that human genomes encoded large numbers of HFGs and, in turn, created genotypic and phenotypic diversities.

To confirm that human HFGs were conserved and widespread, we investigated whether these HFGs existed in the Genotype-Tissue Expression (GTEx) fusion genes. We used SCIF to analyze 427 GTEx blood samples (dbGap-accession: phs000424.v7.p2) and identified 134,090 fusion transcripts. Fig.1f showed that 576 (48.8%) of 1180 HFGs were present in the total fusion genes found in GTEx’s blood samples. On the other hand, Fig.1g showed that 7,989 fusion transcripts were found in both MZ twins and GTEx’s blood samples and accounted for only 8.7% of the total MZ twins’ fusion transcripts. The former was more than fivefold higher than the latter, confirming that the probabilities of these HFGs inherited by the MZ twins were conserved and had significantly higher frequencies in general populations than other fusion transcripts. Supplementary Table 4 showed that these 576 HFGs were present in 420 GTEx blood samples and ranged from 0.2% to 40.1%, while the MZ twins’ counterparts ranged from 1.4% to 67.7%, the average of which was 10.4%. The former average frequency was 1.4% and was sevenfold less than the MZ twins’ counterpart, reflecting genetic differences between the two populations. Supplementary Table 6 showed that 98.4% of 427 GTEx samples had 1 to 37 HFGs, and the average was 7.9 HFGs, supporting the fact that HFGs were conserved, extremely diverse, and widespread.

To understand the potential mechanisms of generating these diverse HFGs, we arbitrarily classified HFGs into five groups: within-a-gene inversion, inversion, deletion, intra-chromosomal fusion genes, and inter-chromosomal fusion genes. Fig.2a showed that *LIMS1-LIMS1* was a within-a-gene inversion, more likely to be generated via direct *LIMS1* gene tandem duplications. Since SCIF had deliberately removed highly repetitive sequences, identifying within-a-gene inversion HFGs was due to gene homologs and pseudogenes. Hence, the numbers of within-a-gene inversion may be significantly underestimated. Fig.2b showed that head-to-tail *MEG8-SNOR114* genes were rearranged into *SNOR114-MEG8* gene structure by inversion to produce a novel non-coding RNA HFG. Fig.2c showed that *PLEKHO1* and *ANP32E* were located on 1q21 opposite strands to form a tail-to-tail structure, and a potential inversion of *the PLEKHO1* gene might generate head-to-tail *ANP32E-PLEKHO1* HFG. Fig.2d showed that a potential deletion of sequences between *TPM4* and *KLF2* might form *TPM4-KLF2* HFG detected in 54.1% of 74 MZ twins. Fig.2e showed that potential intra-chromosomal translocation produced a *RORA-B2M* HFG. Fig.2f showed that a potential inter-chromosomal translocation resulted in the generation of the *OAZ1-SCO2* HFG. Inter-chromosomal alteration generated 660 HFGs, which counted for 55.9% of 1180 HFGs and met theoretical expectations. As shown in Fig.2, the potential mechanisms to generate HFGs were not different from those observed in somatic genomic alterations, suggesting that the generation of HFGs was a traditional genetic event in the germline cells^8^. Therefore, unless it was under natural selection, any potential fusion gene generated by germline structural variants was a potential HFG and had a much higher frequency than its somatic counterpart.

**Figure 2.**
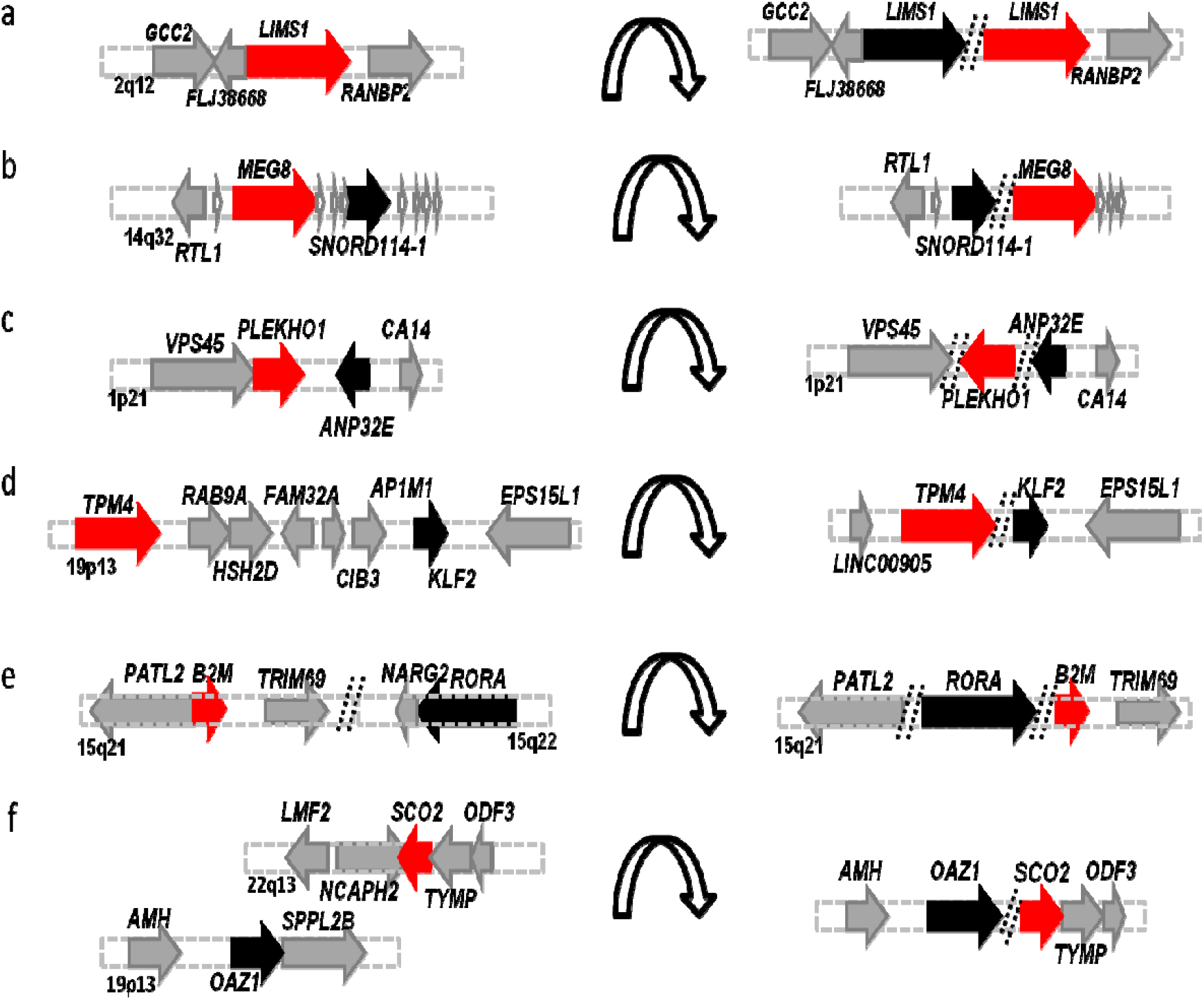
Schematic diagrams of potential genomic alterations for producing human hereditary fusion genes (HGFs). a). Within-a-gene inversion HFG, which was generated via tandem gene duplications. b). *MEG8* was inverted to *SNORD114-1* on the chromosome 14q32. c). Upstream *PLEKHO1* was inverted to downstream of *ANP32E* on the chromosome 1p21 to generate *ANP32E-PLEKHO1*. d) The DNA sequences were deleted between *TPM4* and *KFL2* genes to produce *TPM4*-*KFL2*. e) *B2M* was translocated to downstream of *RORA* on the chromosome 15q21 to generate *RORA-B2M*. f). *SCO2* on the chromosome 22q13 was translocated to downstream of *OAZ1* on the chromosome 19p13 to form *OAZ1-SCO2* fusion gene. Red, black, and gray horizontal arrows represented the 5’ gene, 3’ gene, and genes surrounding both genes, respectively. Horizontal arch arrows showed genomic to produce fusion genes.

To increase SCIF computation speed, we intentionally removed repetitive DNA sequences. To understand the potential roles of repetitive DNAs, we used *the TRNAN35* gene, coding for transfer RNA asparagine 35, as an example to illuminate HFG generation and potential roles during evolution. Fig.3a showed that the *TRNAN35* gene, located at 1q21, was inverted to the positive strand upstream of the *SRGAP2P* gene to form a *TRNAN35-SRGAP2P* HFG. Fig.3b&c showed that *TRNAN35* was translocated to the regions upstream of *FAM91A3P* and *ZNF238* genes to produce *TRNAN35-FAM91A3P* and *TRNAN35-ZNF238* HFGs, respectively. Fig.2d,e,f&g showed that *TRNAN35* was translocated into different chromosomes to yield four putative *TRNAN35*-fused HFGs. Interestingly, *TRNAN35-SRGAP2P, TRNAN35-FAM91A3P, TRNAN35-ACTB, TRNAN35-CHD2*, and *TRNAN35-UBB* were detected in the individual SRR2105730 blood sample. *TRNAN35* could co-regulate the expression of these five HFGs via interactions with aspartyl/glutamyl-tRNA(Asn/Gln) amidotransferase. Hence, they may form a natural network regulated by *TRNAN35*. The addition or deletion of *TRNAN35-fused* HFGs would dramatically increase network diversity and biological diversities. *ALU-SINE* exonization was the extreme example, which increased protein diversity^33,34^ and provided regulatory networks^35^ to human genetic and biological diversity.

**Figure 3.**
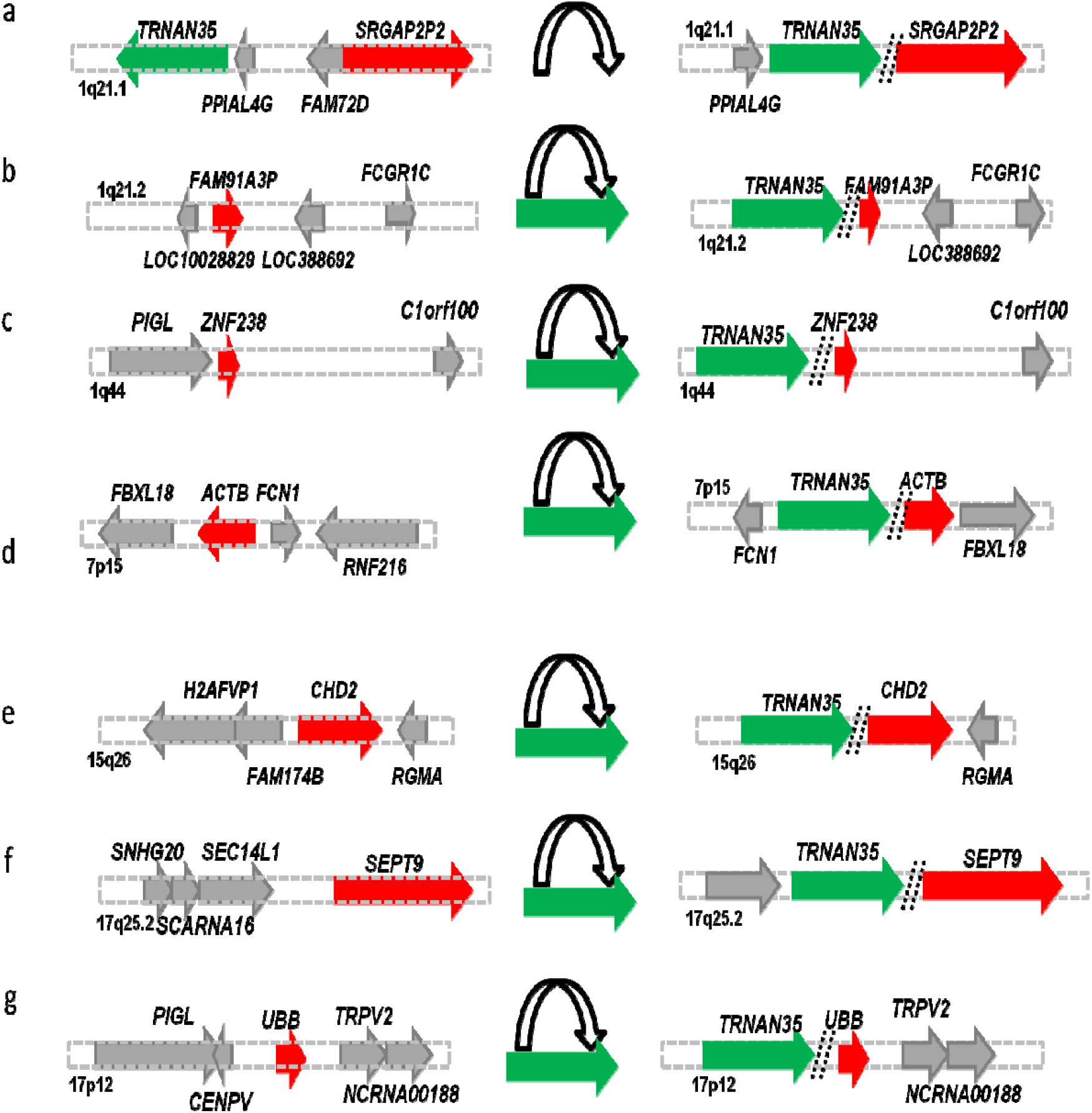
Using the *TRNAN35* gene as an example to demonstrate that repetitive DNA sequences played essential roles in forming hereditary fusion genes (HFGs). a). *TRNAN35* was inverted to upstream of *SRGAP2P* on chromosome 1q21.2 to generate *TRNAN35-SRGAP2P;* b). *TRNAN35* was translocated to upstream of *FAM91A3P* on chromosome 1q21.1 to form *TRNAN35-FAM91A3P;* c). *TRNAN35* was translocated to upstream of *ZNF238* on chromosome 1q44 to yield *TRNAN35-ZNF238;* d). *TRNAN35* was translocated to upstream of *ACTB* on chromosome 7q15 to form *TRNAN35-ACTB;* e). *TRNAN35* was translocated to upstream of *CHD2* on chromosome 15q26 to produce *TRNAN35-CHD2;* f). *TRNAN35* was translocated to upstream of *SEPT9* on chromosome 17q25.2 to form *TRNAN35-SEPT9;* g). *TRNAN35* was translocated to upstream of *UBB* on chromosome 17p12 to generate *TRNAN35-UBB*. Green arrows were *TRNAN35* and 3’ genes. Solid red and gray arrows represented 3’ genes and the genes surrounding *TRNAN35* and 3’ genes, respectively. Horizontal combos of the white arch and solid green arrows showed *TRNAN35* translocations to produce fusion genes. Red and gray arrows represented the genes surrounding 3’ genes and the *TRNAN35* gene.

Family genetic analysis showed that MZ twin inheritance was family-inherited, but no genetic factors have been discovered so far^36^. Hence, we explored whether HFGs were associated with MZ twins’ genetics. Table 1 showed that eight HFGs were detected in over 52.7% of 74 MZ twins’ individuals, ranging from 52.7% to 67.6%, while the GTEx counterparts ranged from zero to 5.2%. The formers were statistically significantly higher than the latter, suggesting that the MZ twins’ inheritance was a complex trait. Half of the eight HFGs, including *LIMS1-LIMS1, SDHAP2-SDHAP2, POM121C-POM121C*, and *PLEKHM1-PLEKHM1* were within-a-gene inversions and originated from tandem duplications (Table 1). The rest were *TPM4-KLF2, BACH1-MECP2, PLXNB2-SCO2*, and *PIM3-SCO2*, the last two of which were *SCO2*-fused HFGs (Fig.4a,b&e) and detected in 67.6% and 52.7% of the MZ twins, respectively. Further analysis showed that *PPP6R2-SCO2* and *TRABD-SCO2* were also from the 22q13 genomic region (Fig.4c&d) and detected in 41.9% and 28.4%, respectively. These four HFGs from the 22q13 region (Fig.4) were significantly higher than *SCO2*-fused HFGs produced via genomic translocations, the highest of which was *OAZ1-SCO2* detected in 24.3% of the MZ twins (Fig.2f). Supplementary Table 6 showed that the recurrent frequencies of these GTEx’s *SCO2*-fused HFGs ranged from 2.1% to 5.2% and were significantly less frequent than the MZ ones, suggesting that *SCO2* amplification and translocation might be involved in the inheritance of MZ twins..The four previously studied cancer fusion genes, *TPM4-KLF2^21^*, *PIM3-SCO2^22,23^, NCO2-UBC^24^*, and *OAZ1-KLF2^21^*, were among the top HFGs associated with the MZ inheritance and detected in 55.6%, 52.7%, 44.6%, and 33.8% of 74 MZ twin siblings, respectively. These suggested that many cancer fusion genes may not be somatic, but HFGs inherited from their parents. This paradigm shift from somatic fusion genes to HFGs will make cancer more predictable, much earlier, easier diagnosed and treated, and more efficient in discovering new therapeutic methods.

**Table 1.** Hereditary fusion genes (HFGs) were significantly associated with the MZ twin inheritance. The GTEx blood samples were used as a control.

| Fusion Gene ID | FT Types | GTEx Blood (427) |  | MZ Twins (74) |  |
| --- | --- | --- | --- | --- | --- |
|  |  | # | % | # | % |
| *PLXNB2-SCO2* | INVERSION | 10 | 2.4 | 50 | 67.6 |
| *LIMS1-LIMS1* | INVERSION | 2 | 0.5 | 50 | 67.6 |
| *SDHAP2-SDHAP2* | INVERSION | 0 | 0.0 | 44 | 59.5 |
| *BACH1-MECP2* | INTER_CHR | 21 | 5.0 | 43 | 58.1 |
| *TPM4-KLF2* | DELETION | 4 | 1.0 | 40 | 54.1 |
| *POM121C-POM121C* | INVERSION | 2 | 0.5 | 40 | 54.1 |
| *PLEKHM1-PLEKHM1* | INVERSION | 2 | 0.5 | 40 | 54.1 |
| *PIM3-SCO2* | INVERSION | 22 | 5.3 | 39 | 52.7 |

The numbers inside the brackets indicated sample sizes.

**Figure 4.**
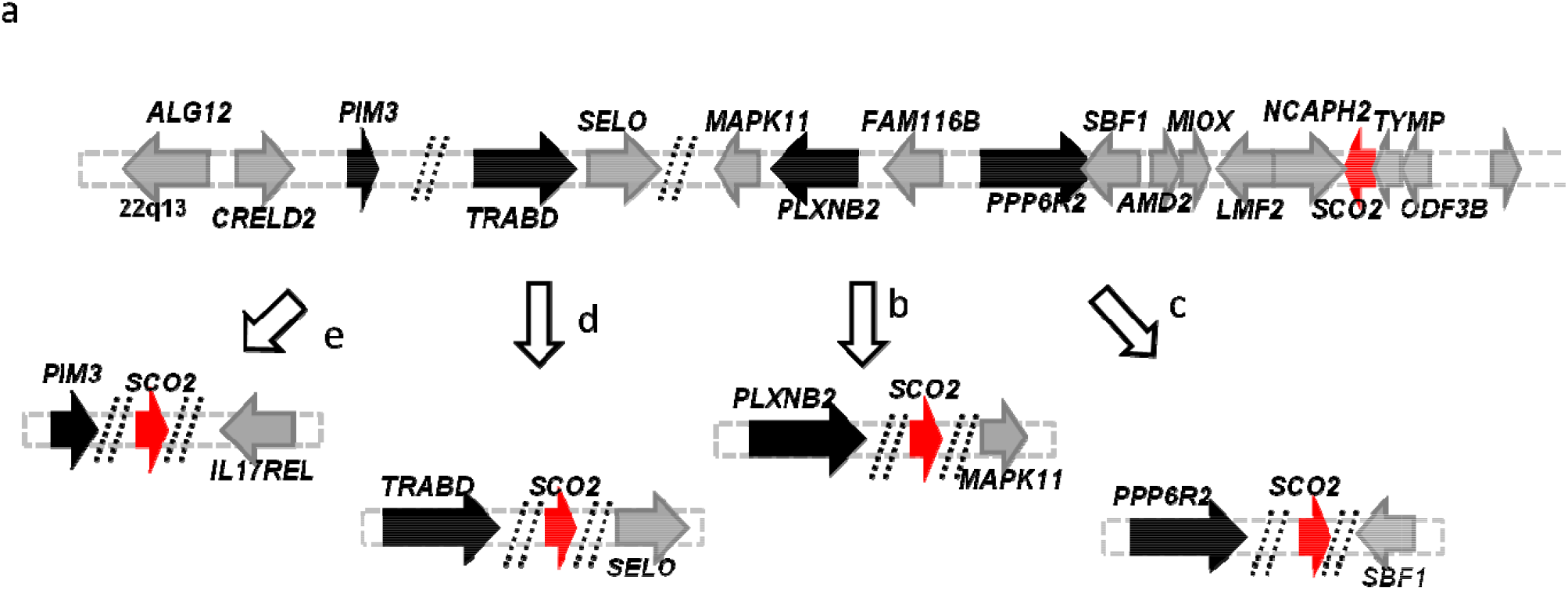
Potential genomic alterations on chromosome 22q13.33 to generate SCO2-fused HFGs. a). The schematic diagram showed the 22q13.33 genomic structure. b). *SCO2* was translocated to downstream of *PLXNB2* to generate *PLXNB2-SCO2*; c) *SCO2* was translocated to downstream of *PPP6R2* to produce *PPP6R2-SCO2;* d) *SCO2* was translocated to downstream of *TRABD* to form *TRABD-SCO2*; e) *SCO2* was translocated to downstream of *PIM3* to generate *PIM3-SCO2*. Solid red arrows were *SCO2* genes. Solid black and gray arrows represented 5’ genes and the genes surrounding the *SCO2* and 5’ genes. White arrows indicated genomic alterations to generate the *SCO2*-fused fusion genes.

In summary, we used MZ twins as a genetic model to identify 1180 HFGs from 37 pairs of MZ twins. The maximum numbers of HFGs were 608 per genome, which were detected in KNAA4 MZ twin siblings. To generate the overlapped HFGs among different groups of MZ twins, we found that MZ genomes encoded over 1000 HFGs. Because a human haploid genome was shown to have 34,234 SVs^10^, a haploid genome might encode thousands of HFGs. As shown in Fig.3, one gene can fuse multiple genes to generate multiple HFGs. Therefore, each gene could generate new fusion genes with every other human gene. If the human genome encoded 25,000 genes^37^, each of which had nine exons^38^, the human genome could generate 5X10^9^ HFGs during evolution. From genomic alterations to HFGs (Fig.2), most of which were due to genomic amplification and duplications (Fig.3), we can predict that offspring could inherit every fusion gene produced via genomic alterations unless it was lethal before the offspring were born. *TPM4-KLF2^21^*, *PIM3-SCO2^22,23^, NCO2-UBC^24^*, and *OAZ1-KLF2^21^*, were among the top HFGs associated with the MZ inheritance, suggesting that many cancer fusion genes may not be somatic but hereditary. If they are authentic, they will lead paradigm shifts in all aspects of cancer studies. These data suggested that HFGs may be the dominant genetic factors associated with many phenotypes and complex traits from MZ twin inheritance to cancer. Recent advances in genome technologies made it possible to directly map genomic SVs and validate HFGs. We can develop more efficient technologies to uncover more HFGs and discover associations between HFGs and diseases and complex traits.

## Methods

### RNA-Seq dataset of monozygotic (MZ) twins

Raw RNA-Seq data of monozygotic (MZ) twins (dbGap-accession: phs000886) was downloaded from NCBI (https://www.ncbi.nlm.nih.gov/Traces/study/?acc=SRP061248). This dataset contained RNA-Seq data from 37 pairs of monozygotic (MZ) twins’ blood samples.

### RNA-Seq dataset of Genotype-Tissue Expression (GTEx)

To evaluate the MZ twins’ hereditary fusion genes, we selected the RNA-Seq dataset of GTEx healthy blood samples as a control. RNA-Seq datasets (dbGap-accession: phs000424.v7.p2) of GTEx’s blood samples were downloaded from NCBI (https://www.ncbi.nlm.nih.gov/projects/gap/cgi-bin/study.cgi?study_id=phs000424.v7.p2). We had identified 427 healthy individual blood samples.

### Identification of fusion transcripts by SCIF (<u>S</u>plicing<u>C</u>odes <u>I</u>dentify Fusion <u>T</u>ranscripts)

SCIF (splicing<u>c</u>odes <u>i</u>dentify <u>f</u>usion transcripts) was described previously by Zhou et al.^26^.

### Classifications of types of fusion transcripts

To better characterize fusion transcripts, fusion transcripts were classified into the following five types based on locations and distances of 5’ and 3’ genes of fusion transcripts.

- **Inter-Chromosomal:** If 5’ and 3’ genes of a fusion transcript were located on two different chromosomes, the fusion transcript was inter-chromosomal.
- **Deletion**: If 5’ and 3’ genes of a fusion transcript originated from the identical chromosomes, the distances between the 5’ and 3’ genes were larger than 1,000,000 bp, and the 5’ and 3’ genes had identical orientations, the fusion transcript was classified as a deletion.
- **Inversion**: If 5’ and 3’ genes of a fusion transcript were mapped to opposite strands of the identical chromosomes or if opposite directions of the same chromosomal stands and the distances between the 5’ and 3’ genes were ≤250,000bp, the fusion transcript was defined as an inversion.
- **Intra-Chromosomal:** If 5’ and 3’ genes of a fusion transcript originated from the identical chromosomes and distances between the 5’ and 3’ genes were longer than 250,000 bp, the fusion transcript was defined as intra-chromosomal.
- **Read-Through:** If 5’ and 3’ genes of a fusion transcript were on the identical chromosomal strands and had the same directions and the distances between the 5’ and 3’ genes were smaller than 250,000 bp, the fusion transcript was classified as a read-through.

### Identification of hereditary fusion genes (HFG)

We specifically defined hereditary fusion genes (HFG) as the fusion genes offspring inherited from parents and excluded epigenetic (read-through) fusion genes. Fusion genes were defined as chimeric genes originating from two different genes whose distances were ≥1,000,000bp. This study used monozygotic (MZ) twins to develop a genetic model to distinguish the somatic fusion genes and hereditary fusion genes. Since MZ twins shared identical genetic materials^29^, If a random SV mutation to generate a fusion gene per individual had a rate of 3.6×10^-2 31,32^, the probability that a pair of MZ twins had the identical SV mutations would be 1.3×10 and was twenty-seven fold less. Therefore, we could use the probability difference to remove a somatic fusion gene. If a fusion gene had been detected in both MZ twins’ individuals (bHFG), this gene had a frequency of 2.7% (1 out of 37), which was 20-fold higher than the random chance of 1.3×10 This difference is statistically significantly higher. If a bHFG had been found in ≥1 pair of MZ twins and this bHFG was found in one individual of another pair of MZ twins (iHFG), the chance of this iHFG that was generated due to a random SV mutation was 3X10^-4^. Therefore, one iHFG was 1/72 or 0.0139, statistically significantly higher than 3X10^-4^. Therefore, this iHFG was counted as a hereditary fusion gene (HFG).

### Identification of epigenetic fusion genes (EFG)

Epigenetic fusion genes (EFG) have been defined as the fusion genes generated via *cis*-splicing of read-through pre-mRNAs of two same-strand neighbor genes. If the distance of the two same-strand neighbor genes were ≤250,000 bp long, the new fusion gene from these two genes was EFG. Since gene orders and the genomic structures were highly conserved in a species or even among different species, the read-through pre-mRNA was due to failed transcriptional terminations and regulated by environmental and physiological factors^20,27,28^. Therefore, we defined the genes to produce the read-through products as EFGs. Healthy individuals had almost identical EFG genomic sequences, and EFGs were frequently detected. EFG expression patterns may be different among different tissues and developmental stages.

## Supplemental Data

**Supplementary Table 1 Gene list of 1180 hereditary fusion genes and their frequencies discovered in each pair of 37 pairs of MZ twins.**

**Supplementary Table 2 The summary information showing individual’s RNA-Seq ID, MZ twin ID, RNA-Seq data size, # of HFGs found in each individuals, and # of HFGs discovered in each pair of MZ twins.**

**Supplementary Table 3 Comparison between KN650 and KNAA4 MZ twins to show that RNA-Seq data qualities affected the numbers of HFGs discovered.**

**Supplementary Table 4 Gene lists and frequencies of 576 HFGs overlapped between the MZ twins and 427 GTEx blood samples.**

**Supplementary Table 5 The numbers of HFGs found for each of 420 GTEx blood samples.**

**Supplementary Table 6 Comparison among the top five SCO2-fused HFGs associated with the MZ twin inheritance.**

## Acknowledgements

We have expressed our most profound appreciation to Ms. Xiaoyan Yang, Prof. Benoit Chabot, Prof. Jeff Xiwu Zhou, Prof. Shunbin Ning, Prof. Yinxiong Li, Dr. Liren Tang, Mr. David Zhuo, and Mr. Noah Zhuo for their various contributions to successfully transform the SplicingCodes theory to the technologies to validate the hereditary and epigenetic fusion genes during the last two decade.

## References

1. Pearson, H. Genetics: what is a gene? Nature 441, 398–401 (2006).

2. Mitelman, F., Johansson, B. & Mertens, F. The impact of translocations and gene fusions on cancer causation. Nat Rev Cancer 7, 233–45 (2007).

3. Jia, Y., Xie, Z. & Li, H. Intergenically Spliced Chimeric RNAs in Cancer. Trends Cancer 2, 475–484 (2016).

4. Wu, H., Li, X. & Li, H. Gene fusions and chimeric RNAs, and their implications in cancer. Genes Dis 6, 385–390 (2019).

5. Mazzarella, R. & Schlessinger, D. Pathological consequences of sequence duplications in the human genome. Genome Res 8, 1007–21 (1998).

6. Puig, M., Casillas, S., Villatoro, S. & Caceres, M. Human inversions and their functional consequences. Brief Funct Genomics 14, 369–79 (2015).

7. Eichler, E.E. Genetic Variation, Comparative Genomics, and the Diagnosis of Disease. N Engl J Med 381, 64–74 (2019).

8. Pocza, T. et al. Germline Structural Variations in Cancer Predisposition Genes. Front Genet 12, 634217 (2021).

9. Huddleston, J. & Eichler, E.E. An Incomplete Understanding of Human Genetic Variation. Genetics 202, 1251–4 (2016).

10. Ho, S.S., Urban, A.E. & Mills, R.E. Structural variation in the sequencing era. Nat Rev Genet (2019).

11. Drummond-Borg, M., Deeb, S.S. & Motulsky, A.G. Molecular patterns of X chromosome-linked color vision genes among 134 men of European ancestry. Proc Natl Acad Sci U S A 86, 983–7 (1989).

12. Reiter, L.T. et al. A recombination hotspot responsible for two inherited peripheral neuropathies is located near a mariner transposon-like element. Nat Genet 12, 288–97 (1996).

13. Hayashi, T., Motulsky, A.G. & Deeb, S.S. Position of a ‘green-red’ hybrid gene in the visual pigment array determines colour-vision phenotype. Nat Genet 22, 90–3 (1999).

14. Bi, W. et al. Reciprocal crossovers and a positional preference for strand exchange in recombination events resulting in deletion or duplication of chromosome 17p11.2. Am J Hum Genet 73, 1302–15 (2003).

15. Higgs, D.R. et al. A review of the molecular genetics of the human alpha-globin gene cluster. Blood 73, 1081–104 (1989).

16. Quinlan, A.R. & Hall, I.M. Characterizing complex structural variation in germline and somatic genomes. Trends Genet 28, 43–53 (2012).

17. Mitelman, F., Johansson, B., Mertens, F., Schyman, T. & Mandahl, N. Cancer chromosome breakpoints cluster in gene-rich genomic regions. Genes Chromosomes Cancer 58, 149–154 (2019).

18. Chase, A. et al. TFG, a target of chromosome translocations in lymphoma and soft tissue tumors, fuses to GPR128 in healthy individuals. Haematologica 95, 20–6 (2010).

19. Babiceanu, M. et al. Recurrent chimeric fusion RNAs in non-cancer tissues and cells. Nucleic Acids Res 44, 2859–72 (2016).

20. Thierry-Mieg, D. & Thierry-Mieg, J. AceView: a comprehensive cDNA-supported gene and transcripts annotation. Genome Biol 7 Suppl 1, S12 1–14 (2006).

21. Roberts, K.G. et al. Genetic alterations activating kinase and cytokine receptor signaling in high-risk acute lymphoblastic leukemia. Cancer Cell 22, 153–66 (2012).

22. Kuhn, J. et al. PIM3-SCO2 Fusion Is a Novel Transcription-Induced Chimera That Is Highly Prevalent In Childhood AML. Blood 122, 2549–2549 (2013).

23. Menezes, J. et al. CSF3R T618I co-occurs with mutations of splicing and epigenetic genes and with a new PIM3 truncated fusion gene in chronic neutrophilic leukemia. Blood Cancer J 3, e158 (2013).

24. Irene J. Locher, W.A., Daniel M. Borràs, M. Willy Honders, Rick H. de Leeuw, Wilma G.M. Kroes, Peter de Knijff, Cornelis A.M. van Bergen, Peter A.C. ‘t Hoen, Szymon M. Kielbasa, Jeroen F.J. Laros, Marieke Griffioen, Hendrik Veelken,. Fusion Transcripts without Corresponding Cytogenetic Abnormalities in Acute Myeloid Leukemia: Implications for AML Pathogenesis,. Blood 130, 2703 (2017).

25. Oliver, G.R. et al. A tailored approach to fusion transcript identification increases diagnosis of rare inherited disease. PLoS One 14, e0223337 (2019).

26. Zhou, J.X. et al. Identification of KANSARL as the first cancer predisposition fusion gene specific to the population of European ancestry origin. Oncotarget 8, 50594–50607 (2017).

27. Ehrlich, J., Sankoff, D. & Nadeau, J.H. Synteny conservation and chromosome rearrangements during mammalian evolution. Genetics 147, 289–96 (1997).

28. Ahituv, N., Prabhakar, S., Poulin, F., Rubin, E.M. & Couronne, O. Mapping cis-regulatory domains in the human genome using multi-species conservation of synteny. Hum Mol Genet 14, 3057–63 (2005).

29. Van Baak, T.E. et al. Epigenetic supersimilarity of monozygotic twin pairs. Genome Biol 19, 2 (2018).

30. Nishioka, M. et al. Identification of somatic mutations in monozygotic twins discordant for psychiatric disorders. NPJ Schizophr 4, 7 (2018).

31. Conrad, D.F. et al. Origins and functional impact of copy number variation in the human genome. Nature 464, 704–12 (2010).

32. Campbell, C.D. & Eichler, E.E. Properties and rates of germline mutations in humans. Trends Genet 29, 575–84 (2013).

33. Krull, M., Brosius, J. & Schmitz, J. Alu-SINE exonization: en route to proteincoding function. Mol Biol Evol 22, 1702–11 (2005).

34. Schwartz, S. et al. Alu exonization events reveal features required for precise recognition of exons by the splicing machinery. PLoS Comput Biol 5, e1000300 (2009).

35. Shen, S. et al. Widespread establishment and regulatory impact of Alu exons in human genes. Proc Natl Acad Sci U S A 108, 2837–42 (2011).

36. Machin, G. Familial monozygotic twinning: a report of seven pedigrees. Am J Med Genet C Semin Med Genet 151C, 152–4 (2009).

37. Pertea, M. et al. CHESS: a new human gene catalog curated from thousands of large-scale RNA sequencing experiments reveals extensive transcriptional noise. Genome Biol 19, 208 (2018).

38. Sakharkar, M.K., Chow, V.T. & Kangueane, P. Distributions of exons and introns in the human genome. In Silico Biol 4, 387–93 (2004).

